# Streptavidin-drug conjugates streamline optimization of antibody-based conditioning for hematopoietic stem cell transplantation

**DOI:** 10.1101/2024.02.12.579199

**Authors:** Aditya R. Yelamali, Ezhilarasi Chendamarai, Julie K. Ritchey, Michael P. Rettig, John F. DiPersio, Stephen P. Persaud

## Abstract

Hematopoietic stem cell transplantation (HSCT) conditioning using antibody-drug conjugates (ADC) is a promising alternative to conventional chemotherapy- and irradiation-based conditioning regimens. The drug payload bound to an ADC is a key contributor to its efficacy and potential toxicities; however, a comparison of HSCT conditioning ADCs produced with different toxic payloads has not been performed. Indeed, ADC optimization studies in general are hampered by the inability to produce and screen multiple combinations of antibody and drug payload in a rapid, cost-effective manner. Herein, we used Click chemistry to covalently conjugate four different small molecule payloads to streptavidin; these streptavidin-drug conjugates can then be joined to any biotinylated antibody to produce stable, indirectly conjugated ADCs. Evaluating CD45-targeted ADCs produced with this system, we found the pyrrolobenzodiazepine (PBD) dimer SGD-1882 was the most effective payload for targeting mouse and human hematopoietic stem cells (HSCs) and acute myeloid leukemia cells. In murine syngeneic HSCT studies, a single dose of CD45-PBD enabled near-complete conversion to donor hematopoiesis. Finally, human CD45-PBD provided significant antitumor benefit in a patient-derived xenograft model of acute myeloid leukemia. As our streptavidin-drug conjugates were generated in-house with readily accessible equipment, reagents, and routine molecular biology techniques, we anticipate this flexible platform will facilitate the evaluation and optimization of ADCs for myriad targeting applications.

## INTRODUCTION

Hematopoietic stem cell transplantation (HSCT) is a lifesaving therapy which provides the best chance to durably treat a variety of hematologic diseases, both malignant and non-malignant. In preparation for HSCT, patients undergo treatment, or “conditioning,” with chemotherapy and/or irradiation to ablate their hematopoietic stem cell (HSC) compartment to enable engraftment of the incoming donor-derived HSCs^1^. For hematologic malignancies, the conditioning regimen also serves to deplete malignant cells that were not killed by the patient’s prior therapies, with more severely myeloablative regimens providing greater antitumor benefit and protection against relapse^2^. However, due in part to the cytotoxicity of conventional conditioning regimens, the benefits of HSCT must be weighed against the risks of treatment-related adverse events, which may be severe enough to prevent older or infirmed patients from accessing the curative potential of transplantation^3^. Furthermore, they may impede the safe application of HSCT for non-malignant blood diseases such as sickle cell anemia^4^.

There has been great interest in leveraging the exquisite specificity of adaptive immune recognition to selectively target and deplete the HSC niche in preparation for HSCT, with the goal of mitigating toxicities from chemotherapy-and irradiation-based conditioning regimens^5^. While some studies have explored the use of cellular immunotherapies for HSC niche clearance^6^, most have focused on conditioning approaches using antibodies and antibody-drug conjugates (ADC). By targeting receptors such as the phosphatase CD45 or the tyrosine kinase c-Kit (CD117), ADC-based regimens have been used in preclinical models to enable HSCT within (syngeneic)^7–9^ and across (allogeneic)^10–13^ histocompatibility barriers with fewer toxicities.

The efficacy of an ADC depends on several factors, but a critical component is the ability of the conjugate to be internalized by target cells so that it can deliver a toxic drug payload able to induce cell death^14^. Consequently, the choice of payload and the chemistry tethering it to the antibody are of paramount importance to ADC biology. For preclinical modeling in the mouse, we and others have utilized the ribosome inactivating protein saporin as a toxic ADC payload^7,8,12,14–16^. Saporin is commercially available in a streptavidin (SAv)-conjugated format, enabling rapid, reliable production of ADCs from any biotinylated antibody^17^. However, our HSCT studies with saporin-based CD45-and cKit-ADCs showed that these conjugates behaved as nonmyeloablative conditioning agents which failed to control tumor burden in the murine A20 lymphoma model^12^. Although nonmyeloablative ADCs are of clear interest for nonmalignant diseases, in which antitumor benefit provided by the conditioning regimen is not necessary, myeloablative conditioning is preferable for acute leukemias as these intensive regimens provide crucial protection against relapse^3^.

We hypothesized that alternative toxic payloads to saporin would yield ADCs endowed with improved myeloablative capacity and antitumor efficacy. To that end, we developed CD45-and cKit-ADCs using pyrrolobenzodiazepine (PBD) dimers as the toxic payload, which in our preliminary studies enabled full conversion to donor hematopoiesis and provided durable protection against an aggressive primary murine acute myeloid leukemia (AML) model^18^. PBD has been successfully utilized in the CD19-targeted ADC loncastuximab tesirine, which is FDA-approved to treat diffuse large B cell lymphoma^19^. Despite this success, toxicities from the highly potent PBD payload remain a potential concern for clinical use, as evidenced by the adverse events leading to discontinuation of the Phase III clinical trial investigating the CD33-targeted ADC vadastuximab talirine as frontline treatment for AML^20^.

Saporin and PBD are just two of many potential ADC payloads, most of which have not been evaluated in ADCs designed for HSCT conditioning. Investigation of alternative payloads may yield ADCs that more optimally balance efficacy with toxicities. Moreover, screening of novel CD45-or cKit-specific antibodies, or antibodies targeting novel receptors, may reveal clones with superior efficacy as HSCT conditioning ADCs regardless of the linked payload. However, direct chemical conjugation of candidate antibodies and drug payloads can become time-consuming, laborious, and expensive, particularly when screening of many antibody-payload combinations is desirable^21^. A system capable of rapidly connecting antibodies to several different drug payloads would greatly facilitate and expedite identification of optimal ADCs for preclinical and translational research. Such a system ideally would also enable cost-effective scale-up of *in vitro* validated ADC candidates for *in vivo* testing in preclinical mouse or non-human primate models.

To address these unmet needs, we used copper-free, strain-promoted azide-alkyne cycloaddition (“Click” chemistry) to construct a novel panel of SAv-drug conjugates^22^. Utilizing a panel of Click-conjugable payloads, we generated ADCs indirectly conjugated to different payloads simply by brief incubation of SAv-drug conjugates with a biotinylated antibody (Figure 1A). Importantly, this platform makes use of readily available reagents and supplies and does not require complex instrumentation for production or purification, making in-house ADC production accessible to any laboratory equipped for routine molecular biology. Herein, we describe the development of this system and its application to identify the most effective payloads for murine and human CD45-ADCs suitable for use as HSCT conditioning and antileukemia agents.

**Figure 1.**
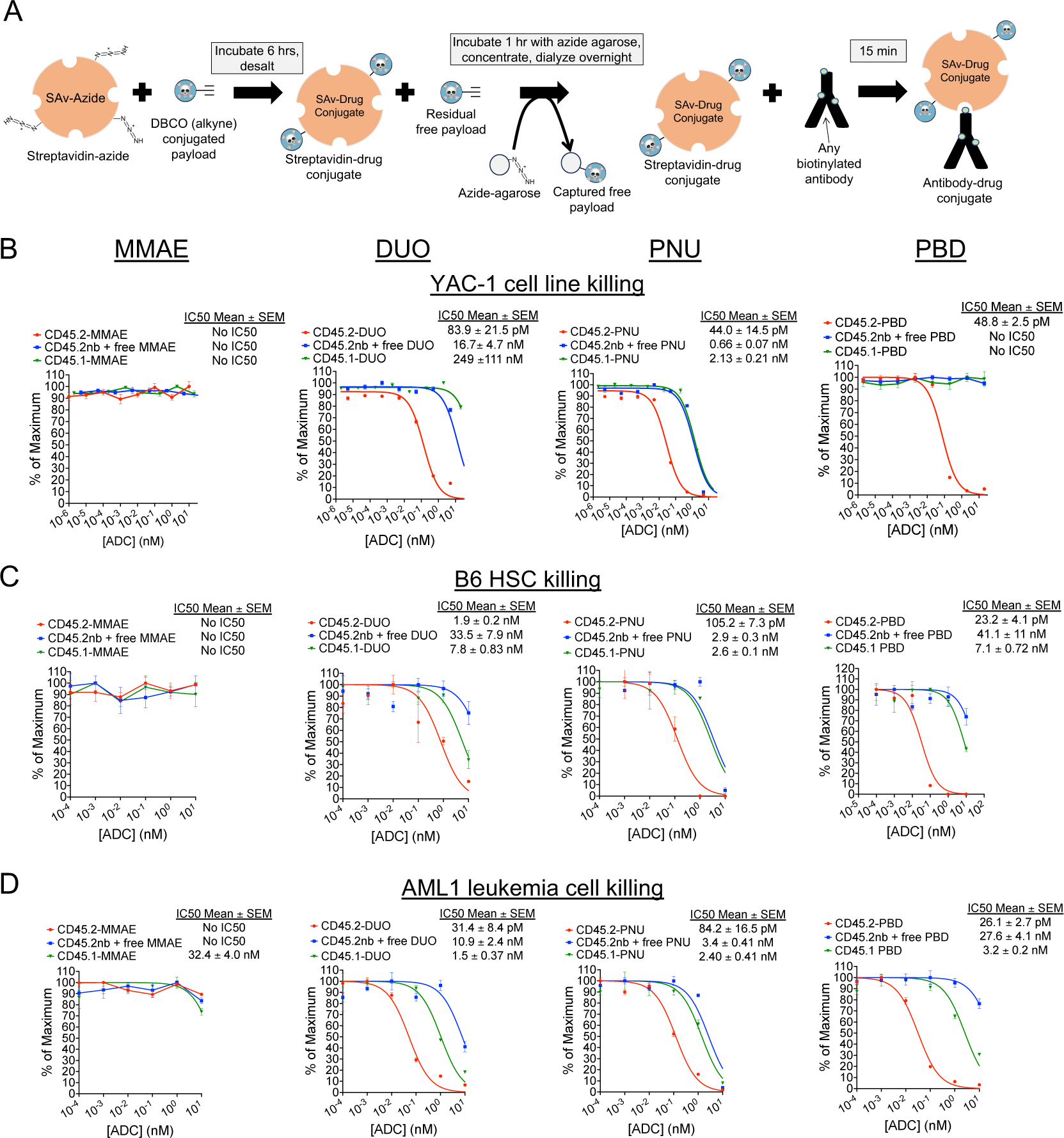
Streptavidin-drug conjugates with DUO, PNU, and PBD payloads yield CD45.2-ADCs that are cytotoxic against murine cell lines, primary HSCs, and AML cells. (A) Schema for production of streptavidin-drug conjugates. (B-D) Cytotoxicity assays of anti-CD45.2-ADC (biotinylated antibody plus streptavidin-drug conjugate), nonbiotinylated (nb) anti-CD45.2 plus free streptavidin-drug conjugate, or CD45.1-ADC (biotinylated isotype control antibody plus streptavidin-drug conjugate) against YAC-1 cells (B), HSCs from B6 mice (C), and AML1 primary leukemia cells (*Dnmt3a^R878H/+^*, FLT3-ITD^+^) (D). Data points represent mean ± SEM of duplicate CFU plates (panel C) or triplicate wells (panels B and D) taken from one representative of at least three independent experiments.

## RESULTS

### *In vitro* evaluation of CD45.2-ADCs produced from SAv-drug conjugates

We selected four drug payloads for initial development and testing of the SAv-drug conjugate platform: the microtubule inhibitor monomethyl auristatin E (MMAE)^23^, the DNA alkylating agent Duocarmycin SA (DUO)^24^, the alkylator and nemorubicin metabolite PNU-159682 (PNU)^25^, and the PBD dimer SGD-1882 (PBD)^26^. Each of these payloads contains cathepsin-cleavable linkers to facilitate intracellular payload release and dibenzylcyclooctyne (DBCO) moieties to undergo the cycloaddition reaction with azide-conjugated protein (Supplemental Figure 1). Commercially prepared SAv-azide was used for these studies to provide a source of conjugation-ready protein with a known number of azide groups per SAv tetramer. Mass spectrometry confirmed successful conjugation of drug payload to SAv, with drug-SAv ratios ranging from 0.55 to 2.5 (Supplemental Figure 2).

We first performed cytotoxicity testing of ADCs made with our SAv-drug conjugates linked to anti-CD45.2 clone 104, which has proven to be a highly effective antibody for ADC-based conditioning in mice^8,12^. As targets for our *in vitro* assays, we used the YAC-1 cell line, whole bone marrow cells from B6 mice (as a source of HSCs), and AML1 primary murine leukemia cells (*Dnmt3a^+/R878H^*/FLT3-ITD^+^; provided by Dr. Timothy Ley)^27^. CD45.2-ADCs conjugated to the DUO, PNU, and PBD payloads were cytotoxic against all target cell types (Figure 1B-D); CD45.2-MMAE was ineffective against all tested target cell types, suggesting MMAE would not be an effective payload for HSCT conditioning in mice. Although CD45-DUO was highly effective against AML1 cells, it showed only modest potency against HSCs, suggesting it too may be suboptimal for HSCT conditioning. CD45.2-PNU and CD45.2-PBD were the most potent at depleting B6 HSCs, with CD45.2-PBD having a wider therapeutic window when compared to our two negative control conditions (nonbiotinylated antibody plus free SAv-drug conjugate or CD45.1-PBD). Thus, *in vitro* screening of CD45-ADCs made possible by our SAv-drug conjugate platform vetted the PNU and PBD payloads as potentially effective for depleting HSCs in preparation for transplant.

In the process of these initial cytotoxicity experiments, we conducted a series of quality control studies to evaluate the robustness of the SAv-drug conjugate system, focusing on SAv-PBD as this conjugate yielded the most effective CD45.2-targeted ADCs in our studies. First, since the purpose of indirect conjugation via the SAv-drug conjugate system is to provide a fast yet effective alternative to direct antibody-payload conjugation, we evaluated whether the cytotoxicity of CD45.2 PBD produced with SAv-PBD was comparable to CD45.2-PBD produced via direct conjugation methods. Indeed, when compared to directly conjugated ADCs made using either a Click chemistry-based method or using the more standard maleimide-thiol coupling chemistry, indirectly conjugated CD45.2-PBD made with SAv-PBD showed comparable levels of cytotoxicity against YAC-1 cells, B6 HSCs, and AML1 cells (Supplemental Figure 3). Next, we compared the activity of CD45.2-PBD conjugates made using four separate batches of SAv-PBD, finding minimal lot-to-lot variability in yields, specific activity, or nonspecific toxicity (Supplemental Figure 4A-B). Finally, since we routinely aliquot and store SAv-drug conjugate stocks at -20°C, we subjected an aliquot of SAv-PBD to repeated freezing and thawing, finding no discernible loss in activity with at least three freeze-thaw cycles (Supplemental Figure 4C).

### *In vivo* testing of CD45.2-ADCs as conditioning agents for murine HSCT

To identify the most effective candidates as conditioning agents for HSCT, we tested the ability of our CD45.2-ADCs to deplete HSCs *in vivo* and evaluated their impact on complete blood counts (CBC) and major peripheral leukocyte subsets. ADC doses of 60 μg were chosen for these studies as this is the molar equivalent of the 75 μg dose of the indirectly conjugated CD45.2-saporin ADCs we used previously^12^. Acute exposure to the ADCs was generally well tolerated, but reduced spleen cellularity and significant weight loss were observed with both CD45.2-and CD45.1-PNU, indicating nonspecific toxicity (Figure 2A and 2B). As expected from our *in vitro* cytotoxicity studies, CD45.2-PNU and CD45.2-PBD effectively depleted mouse phenotypic HSCs (CD150^+^CD48^-^LSK) as well as hematopoietic stem and progenitor cells (HSPCs) in the bone marrow, and markedly reduced colony forming activity (Figure 2C, Supplemental Figure 5). Marrow ablation by the CD45.2-ADCs was largely due to specific toxicity, as CD45.1-PNU and CD45.1-PBD conjugates did not deplete HSCs or HSPCs and only partially reduced colony formation. By contrast, CD45.2-DUO, CD45.2-MMAE, and their respective CD45.1 control conjugates did not markedly affect cell populations in bone marrow, spleen, or peripheral blood. Given these results, CD45-PBD and CD45-PNU proceeded to testing as conditioning agents for HSCT.

**Figure 2.**
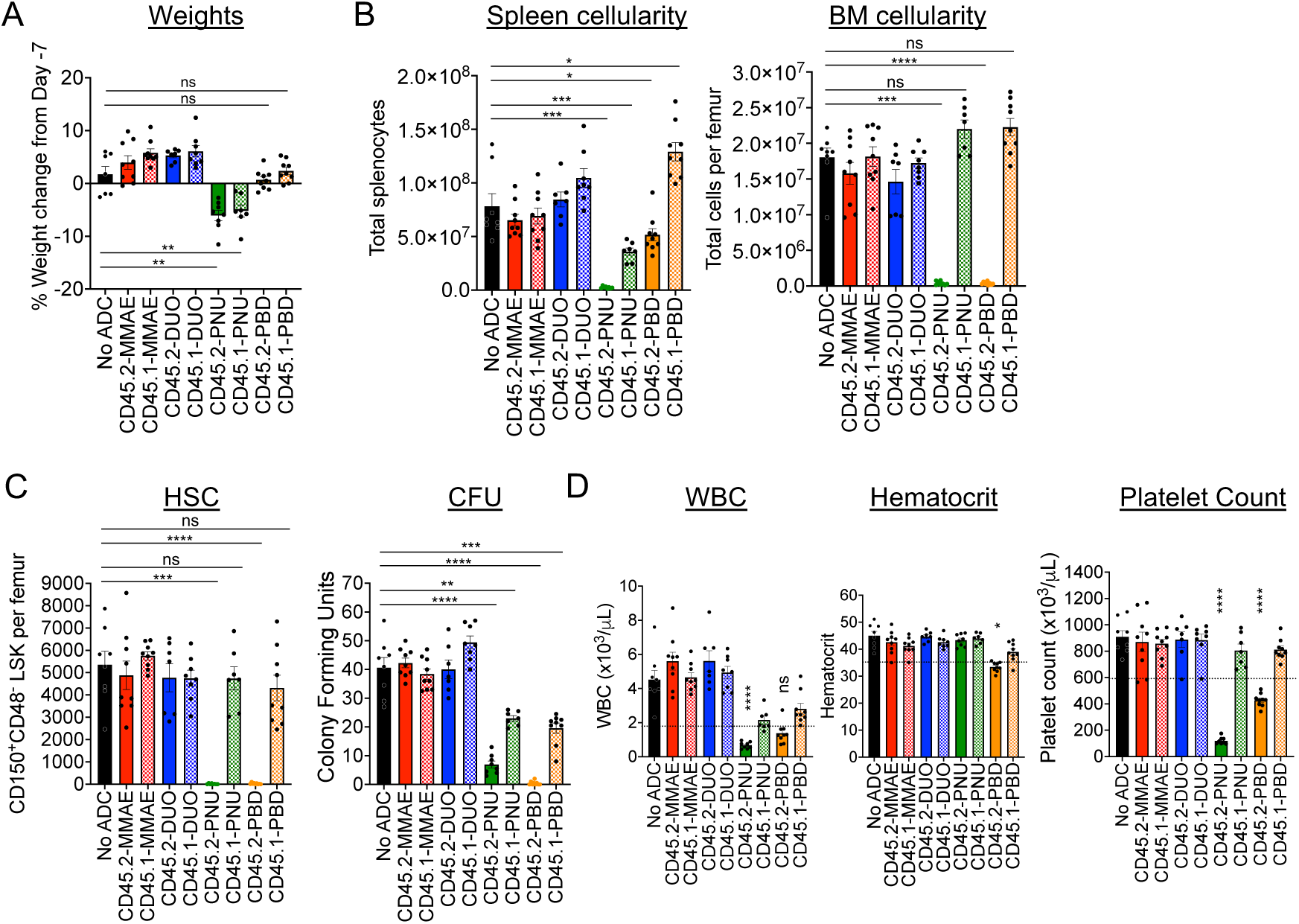
CD45.2-PNU and CD45.2-PBD deplete B6 HSCs *in vivo*, with greater nonspecific toxicities for the PNU payload. B6 mice were either untreated (No ADC) or treated with 60 μg ADC produced by combining the indicated biotinylated antibodies with SAv-drug conjugates. Mice were then sacrificed 7 days later (Day 0) and analyzed. **(A)** Weight change on Day 0 compared to immediately before ADC treatment on Day -7. **(B)** Total cellularity of spleen and bone marrow. **(C)** Counts of CD48^-^CD150^+^ Lin^-^Sca1^+^c-Kit^+^ cells in bone marrow (SLAM-LSK population; HSC), and total CFU from bone marrow of ADC-treated mice assessed after an 8-day incubation in complete mouse methylcellulose. **(D)** Complete blood counts; dotted lines indicate the lower end of the reference range for each cell count. For all panels, each data point represents a single mouse, and bars represent mean ± SEM from mice accumulated over three independent experiments. Statistics: One-way ANOVA with Dunnett’s multiple comparisons test (normally distributed datasets) or Kruskal-Wallis test with Dunn’s multiple comparisons test (non-normally distributed datasets); ns = not significant, * = p < 0.05, ** = p < 0.01, *** = p < 0.001, **** = p < 0.0001.

We employed a syngeneic HSCT model (B6-GFPèB6) to test the ability of CD45.2-PBD and CD45.2-PNU to make marrow space for transplantation (Figure 3A). CD45.2-PBD conditioning was the more effective of the two ADCs, enabling near complete conversion to donor hematopoiesis among myeloid and B cells and high-level mixed chimerism among T cells (Figure 3B). In contrast, despite depleting phenotypic HSCs (CD48^-^CD150^+^ LSK) as well as CD45.2-PBD (Figure 2C), CD45.2-PNU enabled overall donor chimerism of ∼50%. For both ADCs, post-HSCT CBCs were stable and remained within the reference range for the duration of the experiments (Figure 3C), and the levels of engraftment observed in peripheral blood were consistent with those seen among leukocyte subsets in spleen and HSPC subsets in bone marrow (Supplemental Figure 6). Notably, the degree of engraftment enabled by the isotype-matched CD45.1-PBD was considerably lower than the moderately high nonspecific engraftment (up to 40% donor chimerism) we and others have observed previously in mice treated with isotype control antibodies directly conjugated to PBD^13,18^. Taken together, our results show PBD to be our most effective payload for CD45.2-ADC-based HSCT conditioning in the mouse, being both well-tolerated and showing specific ablation of HSCs with minimal depletion of mature hematopoietic cells in the periphery.

**Figure 3.**
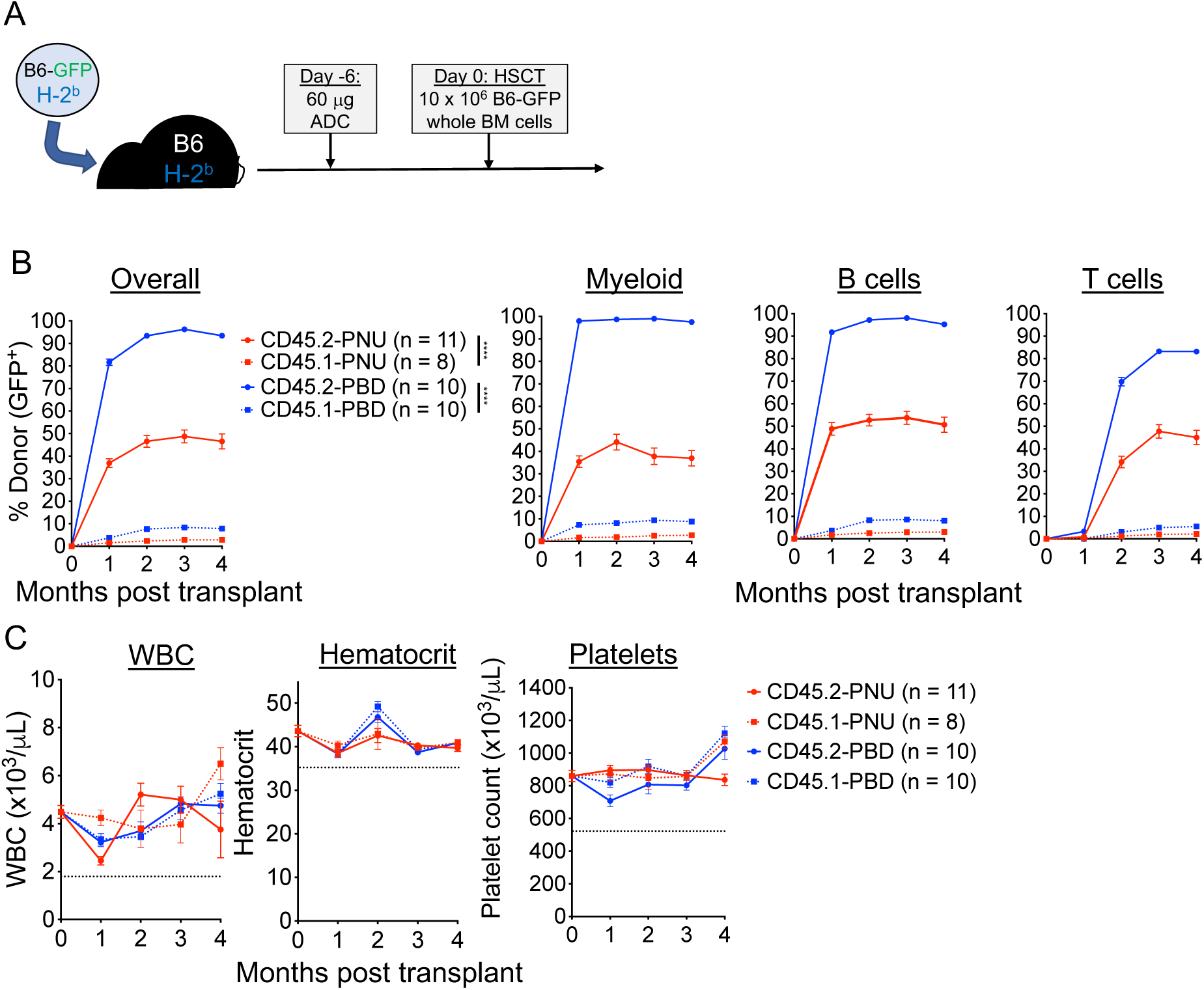
CD45.2-PBD enables syngeneic HSCT with near complete conversion to donor chimerism. **(A)** Schema for syngeneic HSCT of GFP-labeled B6 whole bone marrow into B6 mice, using CD45.2-ADCs made with SAv-drug conjugates for conditioning. **(B and C)** Longitudinal peripheral blood donor chimerism overall and by lineage **(B)** and CBCs **(C)** in recipients conditioned with CD45.2-PNU, CD45.2-PBD, or their respective CD45.1-bound (isotype control) conjugates. Data points indicate mean ± SEM from mice accumulated over two independent experiments. Statistics: Mixed effects model for repeated measures (Panel B); **** = p < 0.0001.

### Evaluation of human CD45-ADCs for targeting HSCs and leukemia cells

Following validation of an ADC targeting strategy in fully murine models, conducting similar studies using human cells and humanized mouse models is an important next step. However, many antibodies to mouse antigens do not cross-react with their human counterparts, and it is possible that the optimal payload for depleting the target cell of interest may differ between mouse and human. Thus, pivoting from mouse to human experiments generally requires that a successful targeting strategy designed in mice be re-optimized for human cells. The SAv-drug conjugate system is well-suited to bridging this gap between murine and human studies, enabling the facile production and evaluation of human ADCs against a candidate target antigen.

To demonstrate this, we generated CD45-ADCs for cytotoxicity experiments by linking SAv-drug conjugates to biotinylated anti-human CD45 antibodies. We began our studies using anti-CD45 clone BC8, which is the antibody used in the radioimmunoconjugate ^131^I-apamistimab currently in Phase III clinical trials for the treatment of AML^28^. CD45-ADCs made with all four payloads yielded similar, picomolar-range IC50 values when targeting Jurkat cells with CD45-ADCs (Figure 4A). Screening of CD45-PBD conjugates made with a panel of different anti-CD45 antibodies demonstrated similar efficacy between clones against Jurkat cells (Supplemental Figure 7). We therefore decided to focus subsequent human cytotoxicity studies on CD45-ADCs using the BC8 antibody, given that this clone has been utilized clinically, can be obtained commercially, and has an available amino acid sequence to enable production of recombinant antibody as needed^29^. In studies assessing the cytotoxicity of CD45-ADC against human HSCs, CD45-PBD was the only ADC tested that showed specific inhibition of colony formation that surpassed the negative controls (Figure 4B).

**Figure 4.**
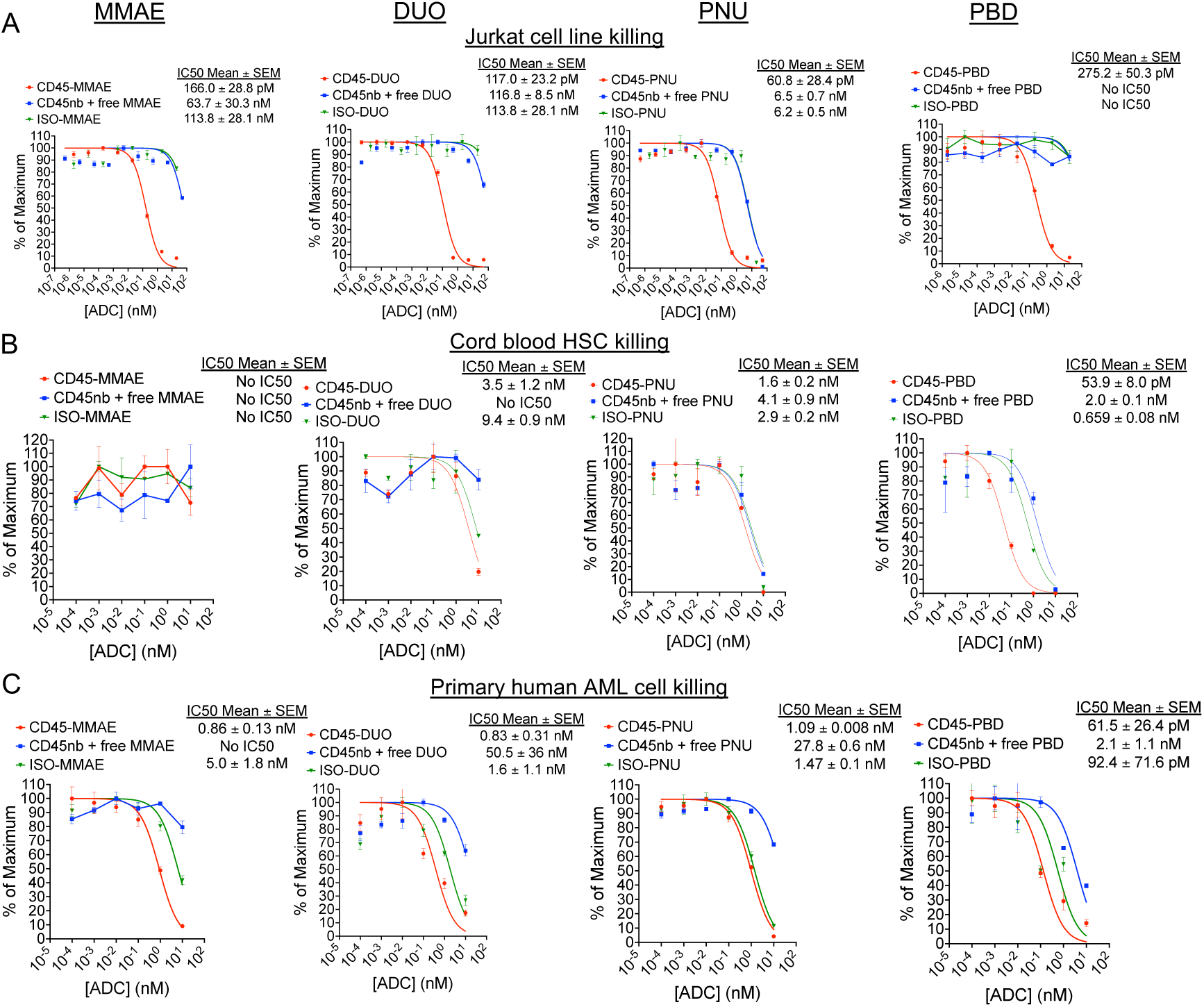
Human CD45-PBD effectively targets and kills human HSC and AML cells *in vitro*. **(A-C)** Cytotoxicity assays of CD45-ADC produced from anti-CD45 clone BC8 (biotinylated antibody plus streptavidin-drug conjugate), nonbiotinylated (nb) BC8 antibody plus free streptavidin-drug conjugate, or mouse IgG1-ADC (biotinylated isotype control antibody plus streptavidin-drug conjugate) against Jurkat cells **(A)**, cord blood derived mononuclear cells **(B)**, and patient-derived leukemia cells **(C)**. Data points represent mean ± SEM of duplicate CFU plates (panel B) or triplicate wells (panels A and C) taken from one representative of at least two independent experiments.

We sought next to examine CD45-ADC cytotoxicity against primary human AML specimens, but the difficulty of culturing primary AML cells *in vitro* is a barrier to this kind of screening^30^. However, we identified a human AML specimen in our departmental biobank that had been successfully passaged and expanded in NSG-SGM3 (NSGS)^31^ mice and which tolerated short-term culture and cytotoxicity testing in complete human methylcellulose medium. Using this approach, we found CD45-ADCs made with all four payloads showed cytotoxicity against human AML, but CD45-PBD showed the highest potency (Figure 4C). Unexpectedly, ADCs with all payloads made with a mouse IgG1 isotype control showed comparable cytotoxicity as the CD45-ADC towards the human AML cells. This degree of toxicity was not observed with human AML cells incubated with nonbiotinylated antibody plus free SAv-drug conjugate, indicating a specific issue with the payload in an antibody-bound form. A possible explanation for this antibody-dependent toxicity is binding and internalization mediated by Fc receptors; notably, CD32 (FcγRII) is highly expressed on our human AML cells, which has moderate binding affinity for mouse IgG1^32^. A role for recognition of and toxicity by mouse IgG1-PBD specifically is suggested by our preliminary observation that two additional PBD-conjugated mouse IgG1 isotype control antibodies were similarly cytotoxic as CD45-PBD, but a mouse IgG2b isotype antibody, to which human FcγRII does not bind^32^, showed less toxicity.

Finally, we tested the ability of CD45-PBD to protect against patient-derived leukemia *in vivo* using humanized mouse models (Figure 5A), using the same human AML cells as were used in our *in vitro* assays (Figure 4C). Since human CD45 antibody clone BC8 does not cross-react with mouse CD45 and does not deplete mouse HSCs when PBD-conjugated, it can be administered to mice without needing to provide stem cell support. CD45-PBD was highly protective in our human AML xenograft model, delaying or preventing leukemia cell expansion over the course of the experiment (Figure 5B-C). While the CD45-PBD treated mice that eventually succumbed to leukemia had clear involvement of peripheral blood, bone marrow, and spleen, surviving mice had no detectable human AML cells in those organs (Figure 5D). Importantly, while the mouse IgG1-PBD conjugate induced significant cytotoxicity against human AML *in vitro*, the same ADC provided minimal survival benefit *in vivo* compared to untreated mice. Thus, our studies demonstrate that CD45-PBD provides specific antileukemia benefit against patient-derived AML *in vivo*.

**Figure 5.**
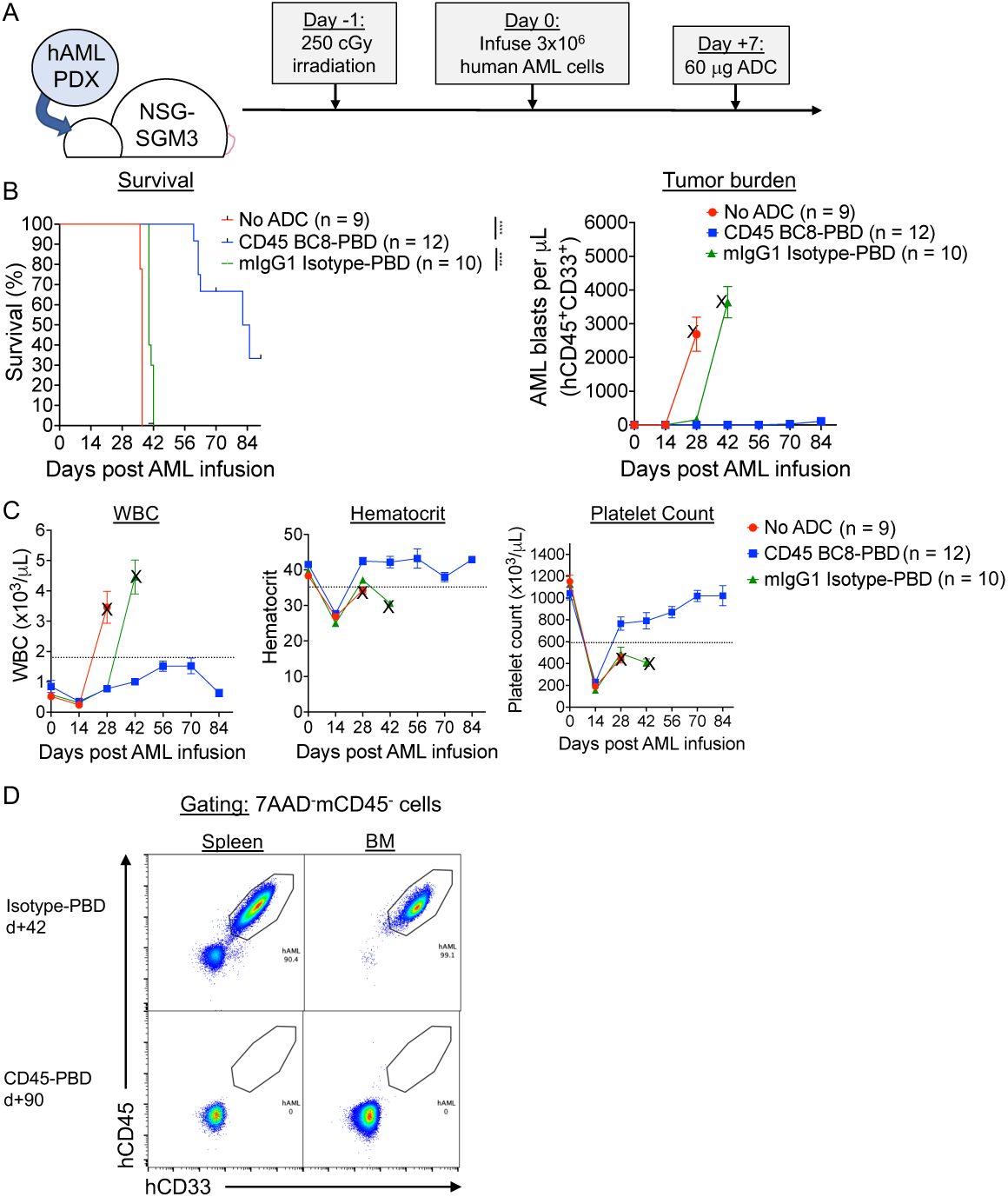
Human CD45-PBD produced from SAv-drug conjugates and anti-CD45 clone BC8 prolongs survival in AML patient-derived xenograft model. (A) Schema for patient-derived xenograft (PDX) AML model in NSGS mice and treatment with CD45-PBD. (B-C) Survival and absolute blast count (human CD45^+^CD33^+^ cells) in peripheral blood (B) and CBCs (C). Death or euthanasia of all individuals in a treatment group is indicated by “X.” (D) Flow cytometry of spleen and BM demonstrating the absence of leukemia cells in surviving CD45-PBD at d+90 versus Isotype-PBD treated mice that succumbed to leukemia at d+42. Plots in panel (D) are gated on 7AAD^-^mCD45^-^ cells. Numerical data are presented as mean ± SEM of mice accumulated over three independent experiments. Statistics: Mantel-Cox log rank test (panel A); **** = *p* < 0.0001

## DISCUSSION

A total of thirteen ADCs are approved globally for clinical use against solid and hematopoietic cancers, with dozens more currently under evaluation in clinical trials^33^. However, many candidate ADCs that showed promising results in preclinical studies did not successfully translate to humans due to inadequate antitumor efficacy and/or unacceptable toxicities. Given that ADCs use payloads that are far too toxic for standalone administration, it is anticipated that even low-level systemic exposure to these compounds would be sufficient to cause adverse effects^34^. Achieving clinical benefit with acceptable toxicities therefore requires careful optimization of the many variables that impact ADC function against a given target cell type, including the antibody, its target antigen, the drug payload, the drug-antibody ratio, the drug conjugation and linker chemistry, and the *in vivo* pharmacokinetic/pharmacodynamic behavior of the conjugate^35^. Methods that facilitate preclinical ADC optimization, encompassing both *in vitro* and *in vivo* modeling, are essential to clinical translation of successful ADC candidates.

In this study, we developed a simple, robust method for conjugating multiple different small molecule ADC payloads to SAv, enabling rapid in-house ADC production starting from a biotinylated antibody or other ligand of interest. The SAv-drug conjugates yielded by this process were of sufficient quantity and quality for both *in vitro* and *in vivo* studies; starting from 2 mg SAv-azide, average yields of ∼1.2 mg conjugated protein were typical which is enough to treat approximately 70 mice with 60 μg ADC (assuming a 1:1 molar ratio of biotinylated antibody to SAv-drug conjugate). We believe this system will greatly streamline development of ADCs for immunotherapeutic purposes, with wide-ranging applications including optimization of existing ADCs, discovery of novel payloads and target antigens, and high-throughput screening of antibody libraries. Furthermore, SAv may also conjugated to novel non-cytotoxic payloads, enabling rapid testing of antibody conjugates with agents such as antibiotics^36^, oligonucleotides^37^, immunomodulators^38^, CRISPR ribonucleoproteins^39^, or proteolysis targeting chimeras (PROTACs)^40^ for therapeutic effect.

Strategies utilizing biotin-SAv coupling for ADC screening and production have been described previously but have limitations which we sought to address via development of our SAv-drug conjugate platform. Commercially available SAv-saporin conjugates, for example, were instrumental to our demonstration that high level donor engraftment could be achieved across immunological barriers without chemotherapy or irradiation-based conditioning^17^. However, we did not observe significant antitumor benefit of CD45-saporin as a single agent in murine lymphoma or AML models^12,18^. Furthermore, the immunogenicity of saporin and its association with adverse events like vascular leak syndrome would limit the clinical utility of saporin, particularly in situations where repeat ADC dosing are necessary^41,42^. The need to evaluate multiple payloads other than saporin then cost-effectively scale up the top candidates for *in vivo* experiments were the main issues for which in-house development of our own SAv-drug conjugates provided a practical solution. A similar strategy using SAv-conjugated antibody with biotinylated payload was recently shown to yield ADCs capable of potent *in vitro* and *in vivo* efficacy^43^. A limitation of that system, however, is that relatively few payloads are commercially available in biotinylated format compared to the conjugation-ready payloads available using amine, thiol, or Click chemistry. Moreover, a major advantage of coupling the payload to SAv rather than biotin is the ability to readily use the wide array of antibodies and recombinant proteins which are either already available in biotinylated format or can be rapidly biotinylated in under one hour with standard conjugation kits.

In our experiments, the PBD dimer SGD-1882 was the most effective payload when used to target HSCs and leukemia cells with mouse and human CD45-ADCs, despite the SAv-PBD conjugate having the lowest drug-SAv conjugation ratio. In contrast to previous work with directly conjugated ADCs^12,13^, minimal HSC depletion and donor engraftment were observed in recipients conditioned with isotype control antibody conjugated to SAv-PBD, suggesting less nonspecific toxicity. The extremely high potency of PBD dimers, coupled with their known myelotoxicity, activity against both dividing and quiescent cells^44^, and capacity for bystander toxicity, may have enabled potent cytotoxicity towards HSCs and AML cells even at low drug conjugation ratios. This, in turn, would allow sufficiently low systemic exposure to PBD to mitigate off-target toxicities. The nonspecific toxicity profile of our SAv-PBD conjugates may also have benefitted from the post-conjugation incubation with azide agarose, which we included to maximize removal of free DBCO-coupled PBD from the final product.

We are currently pursuing several technical optimizations that will address limitations of the current platform and improve its utility in the future. First, although cathepsin-cleavable Val-Ala or Val-Cit linkers are stable in human plasma, they are susceptible to cleavage in mouse plasma via by carboxylesterase 1C (CES1c)^45^, leading to toxicities secondary to premature payload release. This could impact whether a candidate ADC advances to further preclinical or human studies or is excluded from further consideration due to unacceptable toxicities in mouse models. This issue has been circumvented either using mice deficient in CES1c^46^, or by incorporating ADC linkers containing an acidic residue N-terminal to the valine residue which are resistant to CES1c but remain sensitive to cleavage by cathepsins^47^. An alternative approach is the use of non-cleavable linkers^48^, which would improve ADC payload stability in plasma but also hamper specific payload release by requiring target cells to first degrade the antibody intracellularly. However, this may be permissible for a payload like PBD whose extremely high potency could offset the lower cytotoxicity resulting from less efficient intracellular payload release; this would be particularly advantageous if this also minimizes accessibility of free payload to healthy cells. Indeed, proof of principle for non-cleavable PBD-based ADCs was provided by a study showing efficacy of HER2-and CD22-ADCs *in vitro* and *in vivo* in murine breast cancer and lymphoma models, respectively^49^.

A second modification we are pursuing is the use of monomeric SAv molecules for payload conjugation. As we routinely conjugate biotinylated antibody to SAv-drug conjugates at a 1:1 molar ratio, the additional unoccupied biotin binding sites of wild-type tetrameric SAv may not be needed and in fact may promote aggregation by crosslinking biotinylated antibodies via their multiple biotin groups. Normally, the high affinity of wild-type SAv for biotin depends on its tetrameric quaternary structure, with residues from adjacent subunits strongly influencing biotin binding^50^. However, tetrameric SAv with only one intact, wild-type affinity biotin binding site has been described^51^, and monomeric SAv engineered to retain a near-normal k_off_ on biotin binding is commercially available. Beyond minimizing protein aggregation, a particular advantage of monomeric SAv is its smaller size relative to drug payloads; this may enable determination of drug-SAv conjugation ratios via SDS-PAGE rather than mass spectrometry, as was possible for SAv-saporin due to the larger size of saporin (30 kDa) relative to streptavidin (53 kDa).

Finally, an unexplored application of our SAv-drug conjugate system is the generation of dual-payload ADCs. Tumor cell heterogeneity provides an opportunity for the emergence or expansion of treatment-resistant clones, setting the stage for relapsed disease which generally carries a dismal prognosis^52^. Synergistic combinations of ADC payloads with different mechanisms of action can improve antitumor efficacy and reduce the chance for escape variants to develop. Our existing SAv-drug conjugate system could be adapted for use in dual payload ADCs by direct maleimide-thiol conjugation of one payload to a biotinylated antibody, followed by indirect conjugation of the second payload using a SAv-drug conjugate. Alternatively, since SAv natively lacks cysteine residues, recombinant SAv engineered with an N-terminal cysteine and functionalized with azide groups could enable installation of dual payloads on SAv via maleimide-thiol and Click chemistry, respectively. Clear demonstration of benefit for a dual-payload system, however, requires careful consideration of controls, including comparisons with single-payload ADCs (administered alone and together) with normalization of drug-antibody ratios between treatment regimens^53^. The modularity of the SAv-drug conjugate system could facilitate demonstrations of additive or synergistic effects of a dual-payload ADC versus its single payload counterparts.

In summary, improvements in the safety and efficacy of ADCs over the past two decades has led to considerable enthusiasm in their potential as cancer therapeutics. Striking the optimal balance between efficacy and toxicity remains a critical goal in improving the successful translation of ADCs to the clinic. Our study offers tools we believe will enable any laboratory, particularly those embarking on research involving ADC-based targeting, to develop their own novel strategies that will contribute to this ongoing effort.

## ACKNOWLEDGMENTS

This study was funded by NIH/NCI grant R35CA210084 (JFD, including a Research Supplement to Promote Diversity to SPP), NIH/NCI Leukemia SPORE grant (P50CA171963; JFD), NIH/NCI Leukemia SPORE Career Enhancement and Developmental Research Awards (P50CA171063; SPP), an ASTCT New Investigator Award (SPP), an ASH Scholar Award (SPP), awards from Gabrielle’s Angel Foundation for Cancer Research (SPP), and an NCI Research Specialist Award (5R50CA211466; M.P.R). Mass spectrometry analyses were performed by the Mass Spectrometry Technology Access Center at the McDonnell Genome Institute (MTAC@MGI) at Washington University School of Medicine, supported by the Diabetes Research Center/NIH grant P30DK020579, Institute of Clinical and Translational Sciences/NCATS CTSA award UL1TR002345, and Siteman Cancer Center/NCI CCSG grant P30CA091842.

## AUTHOR CONTRIBUTIONS

S.P.P. conceived the research and led the project under the mentorship of J.F.D. A.R.Y., J.K.R., and S.P.P. performed all experiments. E. C. and M.P.R. developed and optimized the human AML xenograft model used in this study. A.R.Y. and S.P.P. wrote the manuscript. All authors read and approved the manuscript prior to submission.

## MATERIALS AND METHODS

### Mice

All animal experiments were done in accordance with a research protocol approved by the Washington University Institutional Animal Care and Use Committee. The following mouse strains were used in this study, either purchased from Jackson Laboratories or bred in-house: C57BL6/J (stock no. 000664), C57BL/6-Tg(UBC-GFP)30Scha/J (B6-GFP; stock no. 004353);

NSG-SGM3 (NSGS; stock no. 013062). All mice were maintained in a specific pathogen-free barrier facility on a 12-hour light/dark cycle with water and standard chow (LabDiet 5053; Lab Supply) provided *ad libitum*. Age-and sex-matched mice 6-12 weeks old were used for all experiments with random assignment of individuals to treatment groups. For retroorbital injections, mice were anesthetized using an isoflurane vaporizer (3% isoflurane in O2 delivered at 1 L/min) and injected with 27 or 29 Ga insulin syringes. For experiments requiring irradiation, a Mark I Model 30 irradiator (^137^Cs source, J.L. Shepherd and Associates,) was used; any mice receiving lethal irradiation and all irradiated NSGS mice were maintained on enrofloxacin in the drinking water beginning 2 days before and ending 2 weeks after irradiation. ADC-conditioned mice did not receive antibiotic prophylaxis.

### Mouse and human cell and tissue preparation

Murine spleens were processed into single-cell suspensions by gently homogenizing tissues with a syringe plunger through a 70-micron filter. Bone marrow was harvested from femurs and tibias by centrifugation (10,000 x g for 15 seconds) using a nested tube method as previously described^54^. Mouse peripheral blood was collected via submandibular bleed using Goldenrod 5 mm animal lancets (MEDIPoint) into K_2_EDTA microtainer tubes (BD). For all specimen types, erythrocyte lysis was done using ammonium chloride-potassium bicarbonate (ACK) lysis, and cell washing and storage was done using PBS + 0.5% BSA + 2 mM EDTA (Running buffer).

Deidentified human umbilical cord blood specimens were obtained from the Cleveland Cord Blood Center. Cord blood units were maintained at room temperature with mild agitation until processing, which occurred within 48 hours of collection. Cord blood mononuclear cells were isolated by Ficoll centrifugation and cryopreserved in 90% FBS/10% DMSO for later use. For CD34^+^ purification, the EasySep Human Cord Blood CD34 Positive Selection Kit II (STEMCELL Technologies) was used; following the manufacturer’s protocol, CD34^+^ purities >90% were routinely obtained.

The murine *Dnmt3a^R878H/+^*/FLT3-ITD^+^ primary AML cells (AML1 cells; GFP^+^) were a kind gift from Dr. Timothy Ley and were generated as described^27^. Splenocytes harvested from AML1 leukemia-bearing mice (>95% leukemia cells based on GFP expression) were cryopreserved without further purification. For patient-derived xenograft leukemia modeling, deidentified primary AML specimens were obtained from a biobank maintained by the Washington University Division of Oncology and expanded via passage into NSGS mice. Leukemia cell stocks were prepared by cryopreservation of splenocytes harvested from leukemia-bearing mice, which were routinely >90% hCD45^+^hCD33^+^.

### Complete blood counts (CBC)

A Hemavet 950 hematology analyzer (Drew Scientific) was used to obtain total white blood cell (WBC) and WBC differential counts, hematocrit, and platelet counts from K_2_EDTA-anticoagulated whole blood. Reference ranges were as follows: WBC 1.8-10.7 x 10^3^ cells/μL, Hct 35.1-45.4%, PLT 592-2972 x 10^3^ cells/μL. CBC data from a cohort of age- and sex-matched B6 recipients (n = 12) was used for the pre-HSCT timepoint (t = 0 months).

#### Cell culture and *in vitro* assays

The YAC-1 and Jurkat cell lines were obtained from ATCC, confirmed to be *Mycoplasma* negative, and maintained in R10 medium (RPMI 1640 (Corning) plus 10% FBS (R&D Systems), 1X GlutaMAX (Gibco), 1X penicillin/streptomycin (Gibco), and 55 μM 2-ME (Gibco)). *In vitro* ADC cytotoxicity against these cell lines was assessed using a 72-hour XTT viability assay per the manufacturer’s recommendations (Cell Signaling Technology) and as described^12^.

For colony forming unit (CFU) assays to measure ADC cytotoxicity against HSCs, whole bone marrow cells from B6 mice or human cord blood mononuclear cells were first prepared in I2 medium (IMDM (Gibco) plus 2% FBS and penicillin/streptomycin) at 2 x 10^6^ cells/mL, then diluted 1:10 into Eppendorf tubes containing concentration series of ADC. Next, 300 μL of each 1:10 dilution mix (containing cells and ADC) was added to 3mL aliquots of complete mouse or human methylcellulose media (R&D Systems; HSC007 and HSC005, respectively) and vortexed thoroughly. Finally, 1.1 mL of this methylcellulose suspension was plated in duplicate 30 mm plates and incubated at 37°C for 8-10 days for mouse HSCs) or 10-12 days for human HSCs.

For ADC cytotoxicity studies against AML cells, murine AML1 cells or human AML cells were prepared in I2 medium at 2 x 10^6^ (mouse) or 5 x 10^6^ cells/mL (human) and diluted 1:10 into Eppendorf tubes containing concentration series of ADC. Next, 50 μL of each dilution mix was added to triplicate wells in 24-well plates, then overlaid with 500 μL of complete mouse or human methylcellulose media and incubated at 37°C for 4 days (mouse) or 3 days (human).

### Production and mass spectrometric analysis of streptavidin-drug conjugates

For production of streptavidin-drug conjugates, streptavidin-azide (SAv-azide; 2-4 azide groups per tetramer, Protein Mods) at 2-5 mg/mL stock concentration in PBS was prepared for payload conjugation by first adding DMSO to 20% final concentration as cosolvent.

Dibenzocyclooctyne (DBCO)-linked ADC toxins (Supplemental Figure 1) were obtained from Levena Biopharma and included the following compounds: DBCO-PEG4-Val-Cit-PAB-monomethyl auristatin E (MMAE; SET0301), DBCO-PEG4-Val-Cit-PAB-Duocarmycin SA (DUO; SET0304), DBCO-PEG4-Val-Cit-PAB-DMAE-PNU159682 (SET0313), and DBCO-PEG4-Val-Ala-pyrrolobenzodiazepine (PBD, specific payload molecule SGD-1882; SET0306). All payloads contained cleavable valine-citrulline or valine-alanine linkers to enable endosomal cathepsin-mediated payload release after ADC internalization. Payloads were prepared for conjugation by first dissolving them in DMSO at 10 mM (200 nmol payload in 20 μL DMSO).

Streptavidin-azide and DBCO-linked payloads were conjugated via a copper-free azide-alkyne cycloaddition (“Click” reaction; Figure 1A). Briefly, 2 mg SAv-azide (37.7 nmol) in 20% DMSO in PBS was mixed with 200 nmol payload at 10 mM in DMSO and incubated 6 hours at 20°C with gentle mixing. After removal of any precipitate by centrifugation (15000 x g, 3 minutes), the supernatant was desalted using Zeba 7 kDa spin columns (ThermoFisher) to remove free payload. As an additional cleanup step, SAv-drug conjugation reactions were next incubated for 1 hour with 100 μL azide agarose slurry with gentle mixing (50 μL packed resin; Click Chemistry Tools/Vector Laboratories), after which the resin was removed using Spin-X columns (Corning). Reaction mixtures were concentrated down to between 0.5-1 mL volume using Amicon 4 mL 10 kDa concentrators (Millipore) then dialyzed using Slide-a-Lyzer MINI 10 kDa dialysis devices (ThermoFisher) per the manufacturer’s recommendations. After dialysis, SAv-drug conjugate concentrations were quantified by BCA assay, resuspended at 1 mg/mL in PBS, sterile filtered with an 0.22 um PES membrane, and stored at -20°C. An Exploris 480 Orbitrap instrument (Thermo Scientific) was used to obtain mass spectra for SAv-drug conjugates, with UniDec peak deconvolution used to determine the drug-SAv conjugation ratio.

#### Production of directly conjugated CD45.2 antibody-drug conjugates

To produce CD45.2-ADCs with drug payloads directly conjugated via Click chemistry, antibodies were first reacted with NHS-azide (ThermoFisher) per the manufacturer’s instructions. Successful attachment of azide groups to antibodies was confirmed by a test conjugation with DBCO-AZDye (Click Chemistry Tools/Vector Laboratories) followed by detection of fluor-conjugated antibodies using UltraComp eBeads (ThermoFisher). Azide-conjugated CD45.2 antibody (4.5 mg each) was then reacted 6 hours with DBCO-linked PBD and purified with the same protocol used for producing SAv-PBD (desalting, azide-agarose incubation, concentration, dialysis, and filter sterilization). To produce CD45.2-ADCs with drug payloads directly conjugated with maleimide-thiol chemistry, 5 mg antibody was reduced with Bond-Breaker TCEP solution (ThermoFisher) for 1 hour at 37°C then desalted with Zeba 7 kDa spin columns to remove TCEP. Reduced antibodies were then reacted 6 hours with MA-PEG4-Val-Ala-PBD (maleimide-linked PBD using SGD-1882 payload; Levena Biopharma SET0212) then purified with the same protocol for SAv-PBD except without an azide-agarose incubation.

#### Conjugation of biotinylated antibodies to streptavidin-drug conjugates

Streptavidin-drug conjugates were mixed in a 1:1 molar ratio with biotinylated antibodies and incubated for 15 minutes at 20°C to produce the indirectly conjugated ADCs used in this study. For conversions between mass and molar concentrations, an approximate molecular weight of 210 kDa was used (150 kDa for IgG plus 60 kDa for SAv-drug conjugates). ADCs were prepared using biotinylated anti-mouse CD45.2 (clone 104; BioLegend), anti-human CD45 (clones T29/33 and BC8 from Leinco Technologies; clones 2D1 and HI30 from BioLegend).

Biotinylated isotype control antibodies (BioLegend) were used to produce control ADCs, including anti-mouse CD45.1 (clone A20; control for anti-mouse CD45.2), mouse IgG2b (clone MG2b-57; control for anti-human CD45 clone T29/33), and mouse IgG1 (clone MOPC-21; control for anti-human CD45 antibody clones HI30, BC8, and 2D1).

After the 15 minute conjugation reaction, ADCs were diluted to the desired concentration in culture medium for *in vitro* assays, or endotoxin-free PBS for *in vivo* administration. All antibodies formulated with sodium azide were exchanged into PBS with Zeba 7 kDa spin columns prior to ADC production. For some *in vitro* experiments, cells were treated with nonbiotinylated antibody plus free SAv-drug conjugate to demonstrate that interaction of antibody with payload was required for cytotoxicity.

### *In vivo* murine HSC depletion and syngeneic HSCT model

For terminal HSC depletion studies, B6 mice were infused with 60 μg ADC via retroorbital injection, then sacrificed 7 days later to assess CBCs, HSC and immune cell depletion from spleen and bone marrow. Body weights were recorded immediately before ADC infusion and upon sacrifice. For syngeneic HSCT experiments, B6 recipient mice were treated with 60 μg ADC then infused 6 days later with 10 x 10^6^ B6-GFP whole bone marrow cells. A 6-day interval between ADC treatment and HSCT was chosen (compared to the 7-day interval in terminal HSC depletion studies) to reduce peritransplant morbidity from cytopenias that developed in mice conditioned with the CD45.2-PNU and CD45.2-PBD conjugates (Figure 2D).

### Human patient derived xenograft model

To assess the antileukemia function of ADCs, 3 x 10^6^ patient-derived leukemia cells (expanded via passage in NSGS mice and cryopreserved) were infused into sublethally irradiated (250 cGy) NSGS mice. Seven days later, recipients received 60 μg CD45- or isotype control ADC and were followed longitudinally for survival, CBCs, and leukemia burden in peripheral blood.

### Data analysis and statistics

All statistical analyses were performed using GraphPad Prism version 10. Normality testing was done with the Shapiro-Wilk test. IC50 values for *in vitro* cytotoxicity studies (XTT viability assays, colony forming assays, AML cell culture in methylcellulose) were obtained by curve fitting the data via nonlinear regression with a three-parameter inhibition model. For comparison of CBC values with the lower reference limit, a one-sample Student’s *t* test was used. For pairwise comparisons of multiple groups against a single control group, a one-way ANOVA with Dunnett’s multiple comparisons test (normally distributed data) or a Kruskal-Wallis test with Dunn’s multiple comparisons test (non-normally distributed data) was used. For pairwise comparisons across three or more groups, a one-way ANOVA with Tukey’s multiple comparisons test (normally distributed data) or the Kruskal-Wallis test with Dunn’s multiple comparisons test (non-normally distributed data) was used. Longitudinal donor chimerism analysis was analyzed using a mixed effects model for repeated measures. Survival was analyzed using a Mantel-Cox log rank test. The criterion for statistical significance for all experiments was *p* < 0.05.

