## Supplemental Table 1 – Supplemental Figures 1-7 for "Streptavidin-drug conjugates streamline optimization of antibody-based conditioning for hematopoietic stem cell transplantation"

**Supplemental Table 1. Flow cytometry antibodies and reagents.**

| Reagent name or target | Species reactivity | Clone | Fluorophore | Vendor(s) |
|---|---|---|---|---|
| CD4 | Mouse | GK1.5 | PE-Cyanine7<br>eFluor 450<br>Brilliant Violet 421 | BioLegend<br>eBioScience<br>BioLegend |
| CD8 | Mouse | 53-6.7 | APC Fire 750<br>Pacific Blue<br>eFluor 450 | BioLegend<br>BioLegend<br>eBioScience |
| CD11b | Mouse | M1/70 | APC | BioLegend |
| CD16/32 | Mouse | 93 | PE-Cyanine7 | BioLegend |
| CD34 | Mouse | SA376A4 | APC | BioLegend |
| CD45 | Human | 2D1 | Biotin | BioLegend |
| CD45 | Human | HI30 | Biotin | BioLegend |
| CD45 | Human | T29/33 | Biotin | Leinco |
| CD45 | Human | BC8 | Purified Functional Grade, In Vivo GOLD<br>Biotin, Functional Grade (Custom) | Leinco<br>Leinco |
| CD45.1 | Mouse | A20 | Biotin<br>Purified | BioLegend<br>BioLegend |
| CD45.2 | Mouse | 104 | Biotin<br>Purified | BioLegend<br>BioLegend |
| CD45R (B220) | Mouse/human | RA3-6B2 | FITC<br>APC<br>eFluor 450<br>Brilliant Violet 605 | BioLegend<br>BioLegend<br>eBioscience<br>BioLegend |
| CD48 | Mouse | HM48-1 | APC | BioLegend |
| CD117 (c-Kit) | Mouse | ACK2 | APC Fire 750 | BioLegend |
| CD150 | Mouse | TC15-12F12.2 | PE-Cyanine7 | BioLegend |
| CD161 (NK1.1) | Mouse | PK136 | PE<br>PE-Cyanine7<br>APC | BioLegend<br>BioLegend<br>BioLegend |
| AZDye 488 DBCO | N/A | N/A | AZDye 488 (structurally equivalent to Alexa Fluor 488) | Click Chemistry Tools (Vector Laboratories) |
| Lineage cocktail (CD3, Gr1, B220, TER-119, CD11b) | Mouse | 17A2, RB6-8C6, RA3-6B2, TER-119, M1-70 | Pacific Blue | BioLegend |
| Ly-6A/E (Sca1) | Mouse | D7 | Brilliant Violet 605 | BioLegend |
| Ly-6G/Ly-6C (Gr1) | Mouse | RB6-8C5 | APC | BioLegend |
| Mouse IgG1k Isotype control | N/A | MOPC-21 | Purified<br>Biotin | BioLegend<br>BioLegend |
| Mouse IgG2bk Isotype control | N/A | MG2b-57 | Purified<br>Biotin | BioLegend<br>BioLegend |
| UltraComp eBeads | N/A | N/A | N/A | eBioscience |
| Viability stains | N/A | N/A | 7-AAD | BD |

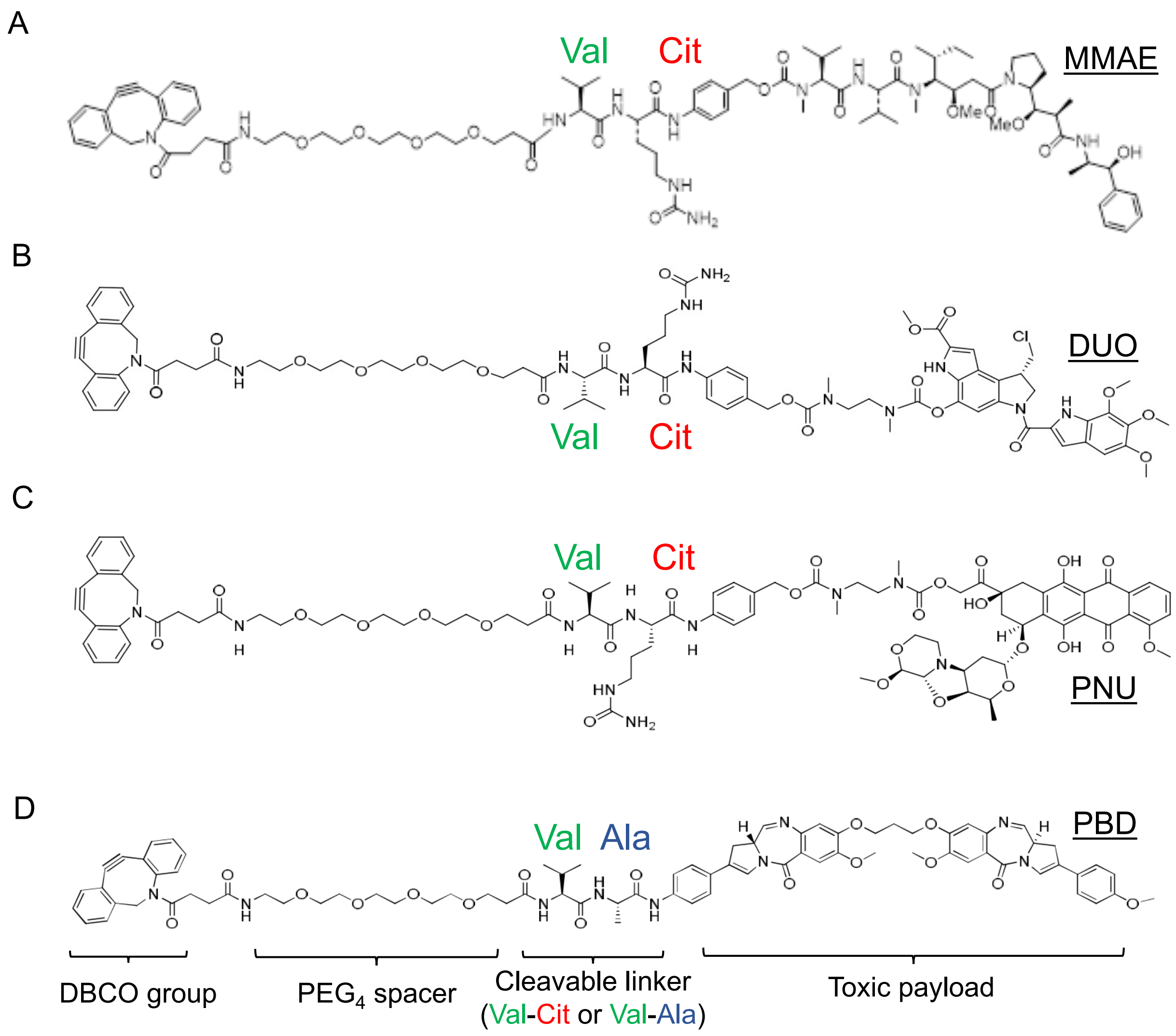


**Supplemental Figure 1. Chemical structures of drug compounds used to produce streptavidin-drug conjugates. (A)** DBCO-PEG4-Val-Cit-PAB-monomethyl auristatin E (MMAE) **(B)** DBCO-PEG4-Val-Cit-PAB-Duocarmycin SA (DUO) **(C)** DBCO-PEG4-Val-Cit-PAB-DMEA-PNU159682 (PNU) **(D)** DBCO-PEG4-Val-Ala-SGD-1882 (PBD). Important structural features are highlighted, including the valine (Val), citrulline (Cit), and alanine (Ala) residues comprising the cathepsin-cleavable linker. Images were provided by the manufacturer.

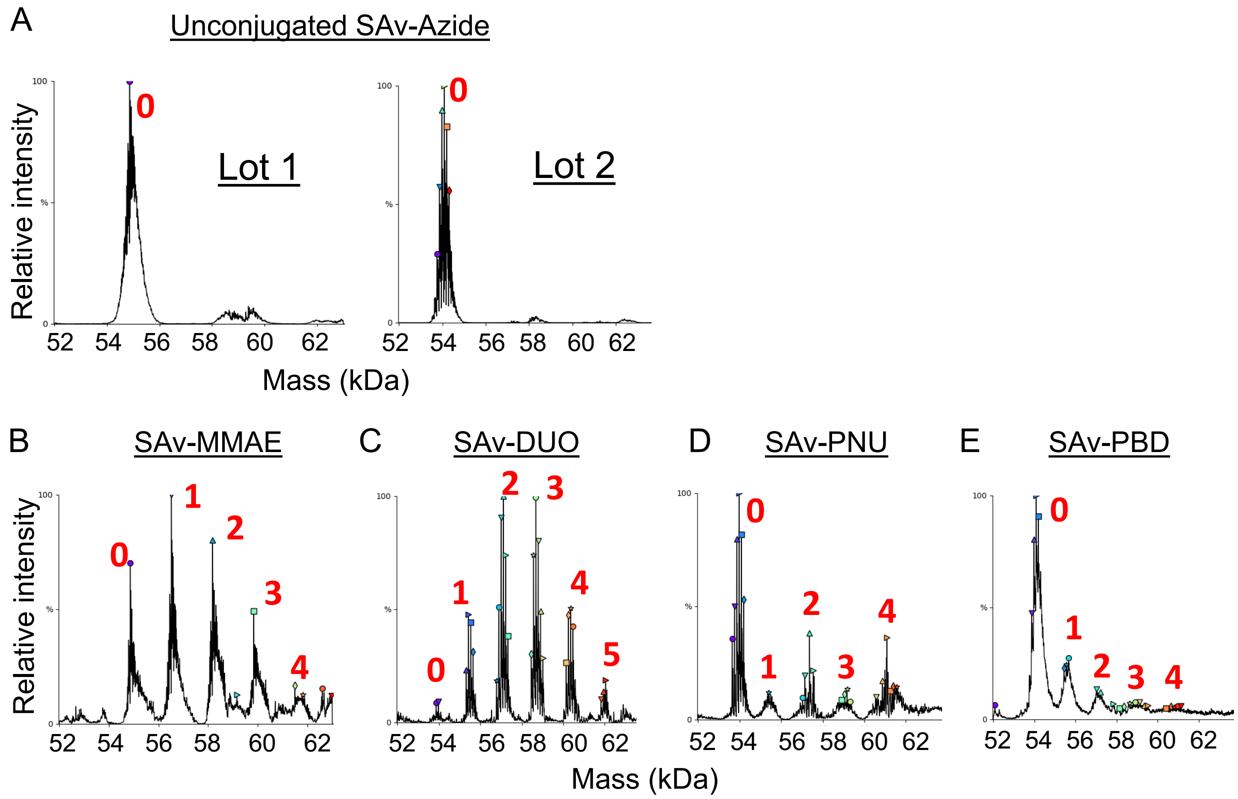


| Replicate | Ratio |
|---|---|
| 1 | 1.53 |
| 2 | 1.55 |
| **Average** | **1.54** |

| Replicate | Ratio |
|---|---|
| 1 | 2.61 |
| 2 | 2.48 |
| **Average** | **2.55** |

| Replicate | Ratio |
|---|---|
| 1 | 1.09 |
| 2 | 1.07 |
| **Average** | **1.08** |

| Replicate | Ratio |
|---|---|
| 1 | 0.5 |
| 2 | 0.6 |
| **Average** | **0.55** |

**Supplemental Figure 2. Mass spectrometry analysis of drug-streptavidin conjugation ratios. (A)** Representative deconvoluted mass spectra of two different lots of unconjugated SAv-azide that were used to produce the streptavidin-drug conjugates that underwent mass spectrometry analysis. Lot 1 is the unconjugated control for SAv-MMAE; Lot 2 is the unconjugated control for SAv-PNU, SAv-DUO, and SAv-PBD **(B-E)** Deconvoluted mass spectra of SAv-MMAE (B), SAv-DUO (C), SAv-PNU **(D)**, and SAv-PBD **(E)** conjugates. The conjugation ratios corresponding to each identified peak are shown in red. All conjugates were tested twice, and the final drug-streptavidin ratios were averages of the ratios obtained from these duplicate runs.

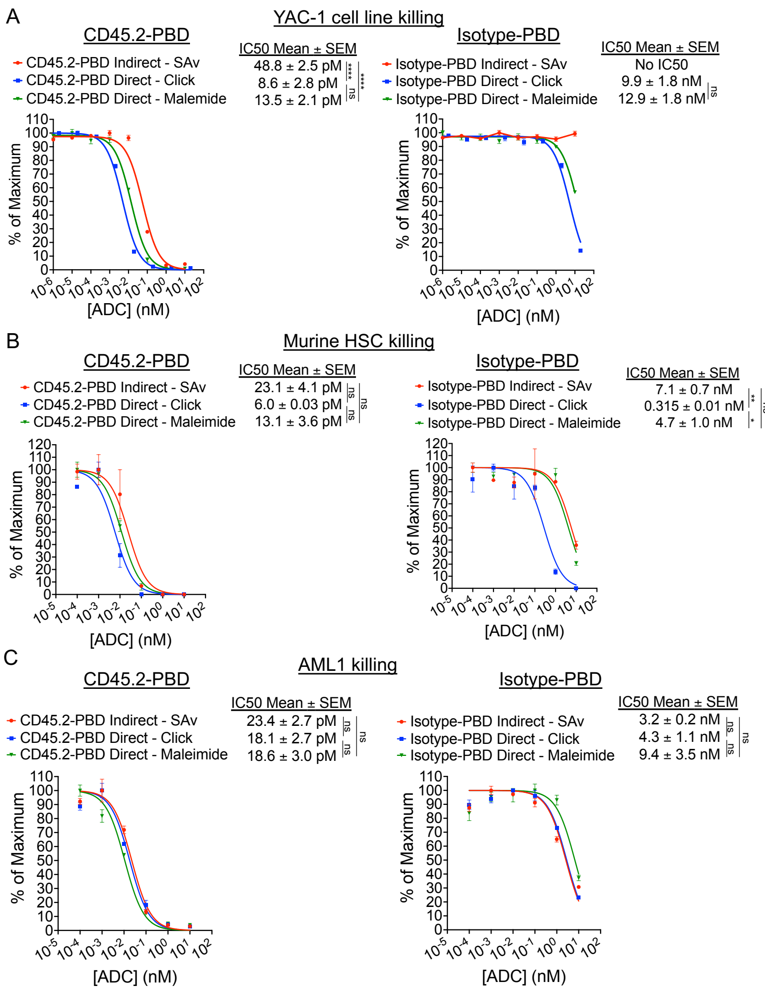


**Supplemental Figure 3. CD45.2-PBD produced with streptavidin-drug conjugates has comparable efficacy to CD45.2-PBD ADCs generated via direct drug conjugation. (A-C)** *In vitro* cytotoxicity of CD45.2-PBD and mouse isotype control-conjugated PBD (Isotype-PBD) against YAC-1 cells **(A)**, murine HSC from B6 mice **(B)**, and AML1 leukemia cells **(C)**. ADCs were generated either via indirect conjugation by combining biotinylated CD45.2 antibodies with SAv-PBD, direct PBD conjugation to azide-coupled CD45.2 antibody via Click reaction, or direct PBD conjugation to TCEP-reduced CD45.2 antibody via maleimide-thiol chemistry. Data points represent mean ± SEM of duplicate CFU plates (panel B) or triplicate wells (panels A and C) taken from one representative of two or three experiments. Statistics: One-way ANOVA with Tukey's multiple comparisons test (normally distributed datasets) or Kruskal-Wallis test with Dunn's multiple comparisons test (non-normally distributed datasets); ns = not significant, * = $p < 0.05$, ** = $p < 0.01$, **** = $p < 0.0001$

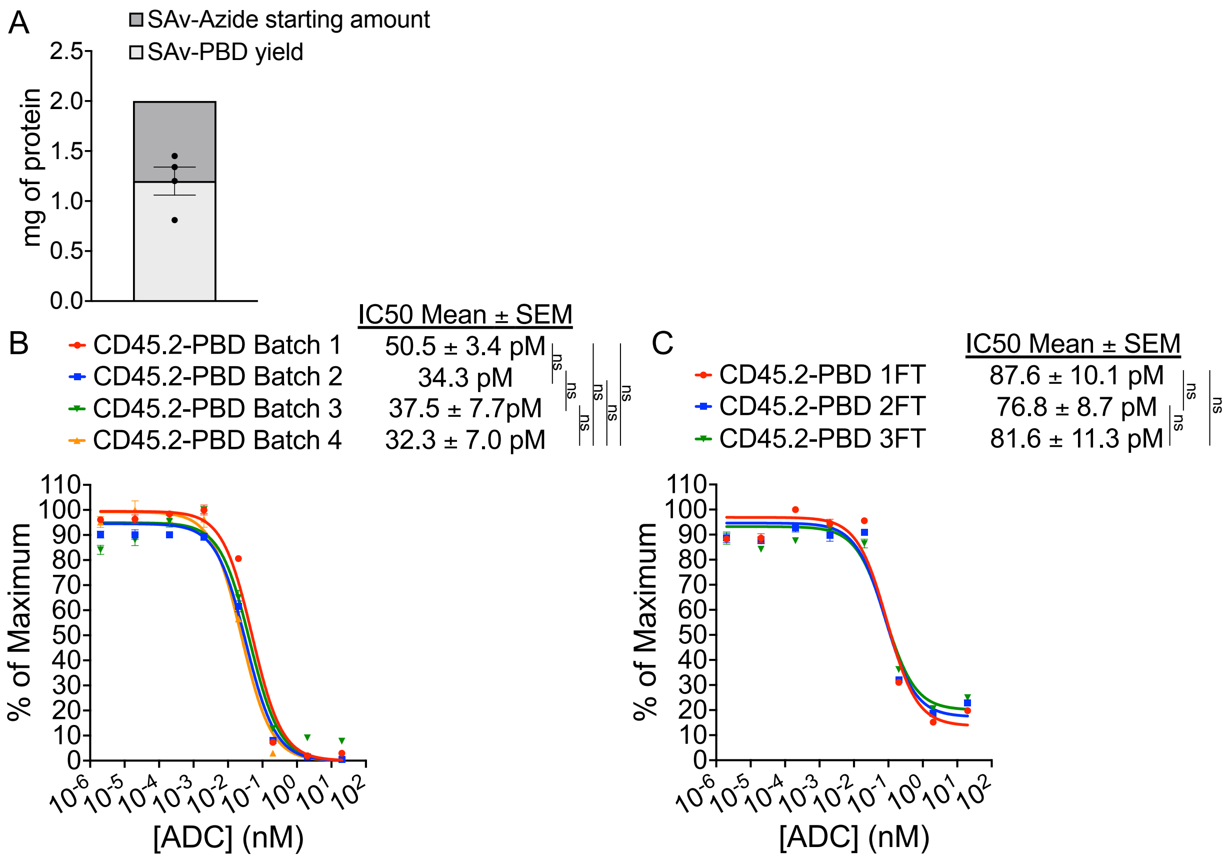


**Supplemental Figure 4. Batch-to-batch variability in conjugate yield and impact of multiple freeze-thaw cycles on cytotoxicity. (A)** Protein yields from four preparations of SAv-PBD quantified by BCA assay. **(B)** YAC-1 cytotoxicity assay performed with CD45.2-PBD made using four separate batches of SAv-PBD with the same lot of biotinylated CD45.2 antibody. **(C)** Three samples of SAv-PBD were taken from a single, never-frozen batch of conjugate and subjected to one, two, or three freeze-thaw (FT) cycles at -20ºC. Each SAv-PBD sample was then combined with biotinylated anti-CD45.2 to produce CD45.2-PBD and tested for cytotoxicity against YAC-1 cells. Data points represent mean ± SEM absorbance at 450 nm from triplicate wells of XTT viability assays that were normalized to the maximum response. IC50 values were obtained from one experiment (CD45.2-PBD Batch 2) or three to four experiments (all other groups). Statistics: One-way ANOVA with Tukey's multiple comparisons test; ns = not significant

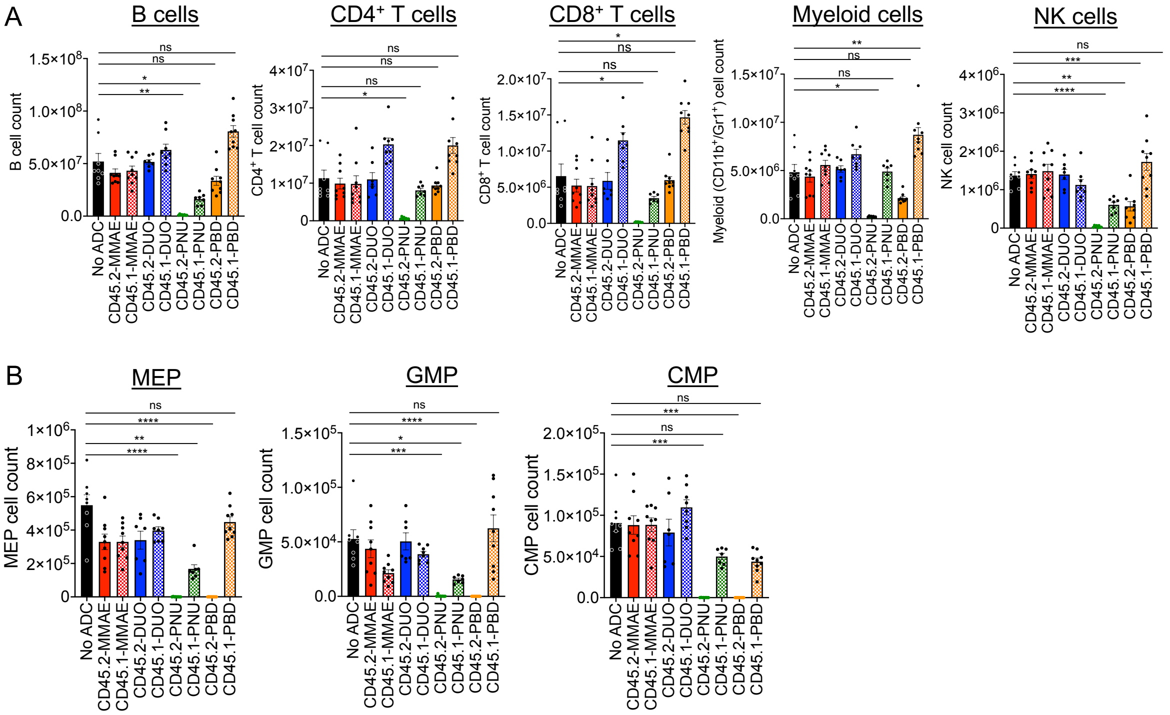


**Supplemental Figure 5. *In vivo* depletion of peripheral leukocyte subsets and hematopoietic stem and progenitor populations by CD45.2-ADCs.** B6 mice were either untreated (No ADC) or treated with ADCs produced by combining the indicated biotinylated antibodies with streptavidin-drug conjugates. Mice were then sacrificed and analyzed 7 days post-ADC treatment as per Figure 2. **(A)** Cell counts of major leukocyte subsets. **(B)** Cell counts of hematopoietic stem cell and progenitor (HSPC) populations in bone marrow; all HSPC populations were identified by CD16/32 and CD34 staining after gating on viable $Lin^{-}Sca1^{-}c\text{-}Kit^{+}$ cells (LK population). Each data point represents a single mouse, and bars represent mean ± SEM from mice accumulated over three independent experiments. Statistics: One-way ANOVA with Dunnett's multiple comparisons test (normally distributed datasets) or Kruskal-Wallis test with Dunn's multiple comparisons test (non-normally distributed datasets); ns = not significant, * = p < 0.05,  ** = p < 0.01, *** = p < 0.001, **** = p < 0.0001.

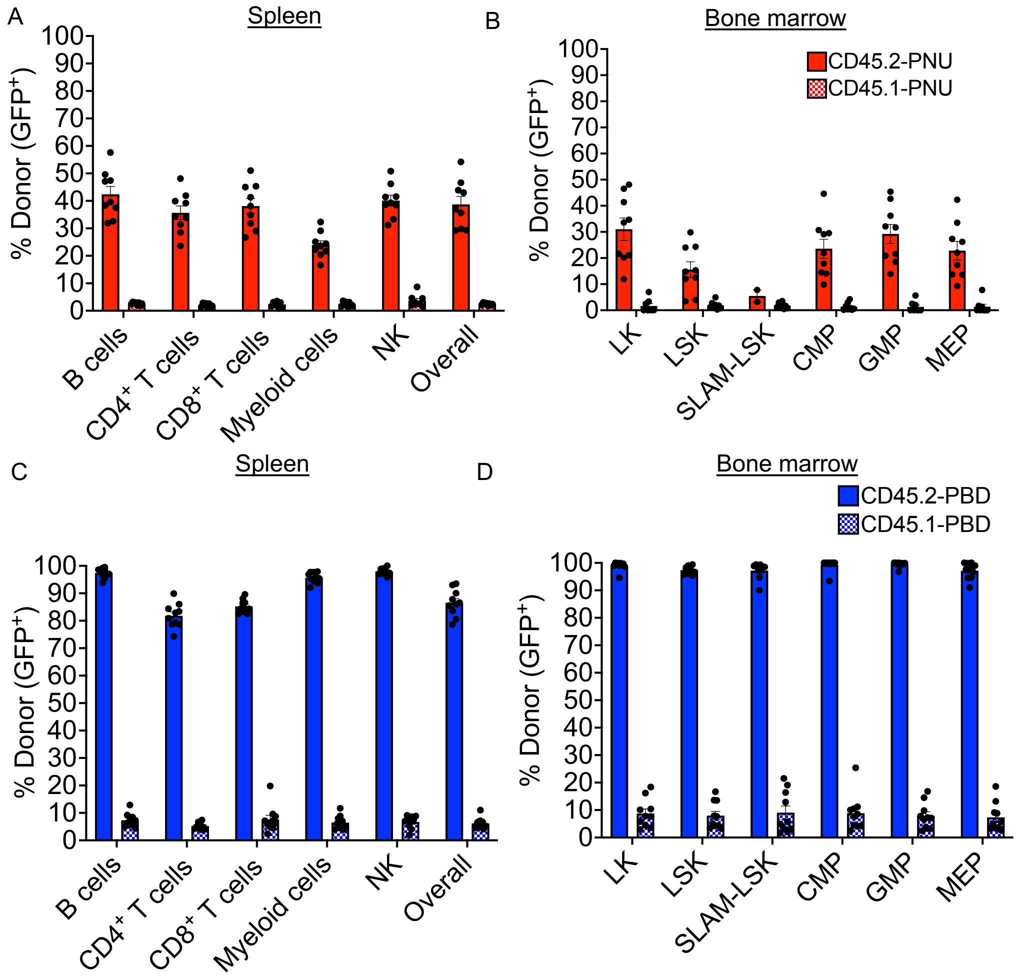


**Supplemental Figure 6. Donor engraftment in spleen and bone marrow of mice conditioned with CD45.2-PBD and CD45.2-PNU. (A-B)** Donor chimerism (frequency of GFP$^+$ cells) overall and by lineage in spleen **(A)** and bone marrow **(B)** in syngeneic HSCT recipients conditioned with CD45.2-PNU performed as per Figure 3. **(C-D)** Donor chimerism overall and by lineage in spleen **(C)** and bone marrow **(D)** in syngeneic HSCT recipients conditioned with CD45.2-PBD. Each data point represents a single mouse, and bars represent mean ± SEM from mice accumulated over two independent experiments.

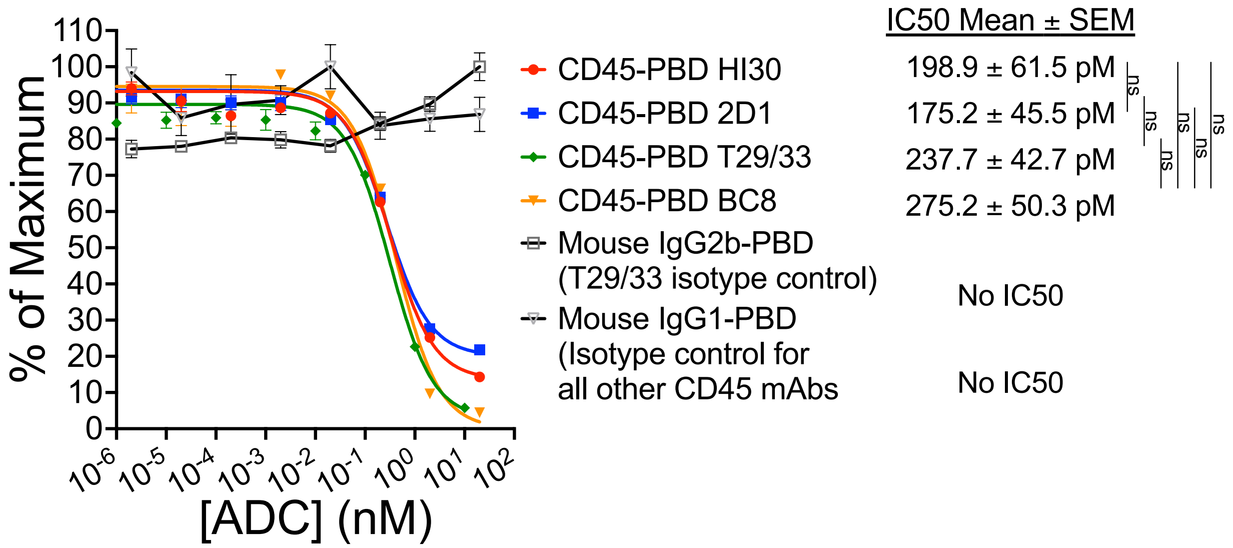


**Supplemental Figure 7. CD45-ADCs produced with different human CD45 antibody clones conjugated to SAv-PBD have similar cytotoxicity against Jurkat cells.** Data points represent mean ± SEM absorbance at 450 nm from triplicate wells of XTT viability assays that were normalized to the maximum response. Mean IC50 values were obtained over four independent experiments. Statistics: One-way ANOVA with Tukey's multiple comparisons test; ns = not significant.
